# Sex in the wild: how and why field-based studies contribute to solving the problem of sex

**DOI:** 10.1101/235465

**Authors:** Maurine Neiman, Patrick G. Meirmans, Tanja Schwander, Stephanie Meirmans

**Affiliations:** Department of Biology, University of Iowa, Iowa City, IA, 52242, USA; Institute for Biodiversity and Ecosystem Dynamics (IBED), University of Amsterdam, P.O. Box 94248, 1090GE Amsterdam; Department of Ecology and Evolution, University of Lausanne, CH-1015 Lausanne; Academic Medical Center (AMC), Universityof Amsterdam, 1105 AZ Amsterdam, NL

**Keywords:** sexual reproduction, asexual reproduction, parthenogenesis, Red Queen, niche differentiation, Muller’s ratchet

## Abstract

Why and how sexual reproduction is maintained in natural populations, the so-called “queen of problems”, is a key unanswered question in evolutionary biology. Recent efforts to solve the problem of sex have often emphasized results generated from laboratory settings. Here, we use a survey of representative “sex in the wild” literature to review and synthesize the outcomes of empirical studies focused on natural populations. Especially notable results included relatively strong support for mechanisms involving niche differentiation and a near absence of attention to adaptive evolution. Support for a major role of parasites is largely confined to a single study system, and only three systems contribute most of the support for mutation accumulation hypotheses. This evidence for taxon specificity suggests that outcomes of particular studies should not be more broadly extrapolated without extreme caution. We conclude by suggesting steps forward, highlighting tests of niche differentiation mechanisms in both lab and nature and empirical evaluation of adaptive evolution-focused hypotheses in the wild. We also emphasize the value of leveraging the growing body of genomic resources for non-model taxa to address whether the clearance of harmful mutations and spread of beneficial variants in natural populations proceeds as expected under various hypotheses for sex.

**Author contributions:** SM and MN conceived the paper idea, SM, PM, MN, and TS designed the review strategy, reviewed and analysed the literature, and wrote the manuscript. All authors gave final approval for publication.

**Data archival location:** The results of our literature survey are provided as electronic supplementary material.

## Introduction

Ask evolutionary biologists about unresolved problems in evolution, and many will question why so many eukaryotic species produce offspring via sexual reproduction (“sex”). The predominance of sex was first identified as a major unanswered question by leading twentieth century evolutionary biologists such as George Williams and John Maynard Smith, who developed theory demonstrating that sexual reproduction should be at a substantial disadvantage vis-à-vis asexual reproduction and thus be rapidly replaced by the latter (Maynard Smith 1971, 1978, Williams 1975). This theory is based on the recognition that sex can impose a variety of costs (e.g., males, recombination, for a recent overview see Lehtonen et al. 2012, Meirmans et al. 2012) that should translate into major advantages for asexual reproduction. The central place of the problem of sexual reproduction in evolutionary theory is illustrated by Graham Bell’s 1982 statement that the maintenance of sex is the “queen of problems” in evolutionary biology (Bell 1982). Within a decade or so after the paradox of sexual reproduction was first identified, dozens of hypotheses for sex had been proposed (Kondrashov 1993). Most of these hypotheses focused on direct or indirect benefits of sexual reproduction that can outweigh (at least in principle) the costs of sex (reviewed in Neiman and Schwander 2011) and that are linked to genetic consequences of meiotic recombination and segregation (e.g., Agrawal 2009a,b). Despite all of this attention and the forty or so years that have passed since the problem of sex was identified, the evolutionary mechanisms underlying the maintenance of sex are still unclear (e.g., Sharp and Otto 2016, Neiman et al. 2017).

The crux of the problem of the maintenance of sex is the persistence of sexual reproduction in so many species, and especially the maintenance of sex in natural populations of organisms or lineages that can use asexual reproduction to produce offspring. Even so, most recent attention towards empirical tests of mechanisms favoring sex has focused on results generated by experimental evolution in laboratory settings. Laboratory studies, which nearly always use genetic model systems (e.g., *Drosophila melanogaster* (Singh et al. 2015), *Saccharomyces cerevisiae* (McDonald et al. 2016), *Tribolium castaneum* (Lumley et al. 2015), *Brachionus calyciflorus* (Becks and Agrawal 2012)), are very powerful because the focal mechanisms can be manipulated and isolated. Indeed, lab-focused studies have provided important tests of the potential for particular mechanisms for sex to be applicable under specific circumstances (e.g., sexual selection-facilitated clearance of mutational load, strong selection for adaptation to new environments; reviewed in Sharp and Otto 2016).

A critical complement to these insights from the laboratory will come from characterization of the mechanisms contributing to the maintenance of sex in natural populations. In particular, despite the rigor and elegance of many laboratory studies, it can be difficult to determine whether their outcomes can be extrapolated to natural conditions. This challenge is exemplified by a recent example of a case where field and lab studies addressing the same mechanisms for sex generated opposite results (Lavanchy et al. 2016).

There are multiple and non-mutually exclusive potential explanations for differences between the outcomes of laboratory and field studies. One important consideration is that model organisms are characterized by short generation times, large populations, and small body sizes, which is not representative of the bulk of metazoans. Model organisms are also often the products of decades of laboratory culture and thus are almost certainly adapted to laboratory conditions (e.g., Sterken et al. 2015). It is also impossible to perform laboratory-based studies featuring all factors that are likely relevant to the realized costs and benefits of sexual reproduction, including locally adapted parasites and predators, competitors for limited resources, extreme abiotic conditions, and unpredictable environmental changes. An additional challenge is that the inclusion of these factors in laboratory experiments can translate into conditions that are unlikely to characterize natural populations, e.g., unrealistically high doses of infectious parasites or the imposition of direct competition between sexual and asexual individuals that experience niche differentiation in the wild. While studies focused on natural populations, and in particular, field-based studies, are more realistic, they cannot match lab studies with respect to the ability to control variables and achieve causal inference. The take-home message is that the insights from field and laboratory studies ultimately need to be integrated in order to produce robust insights into natural patterns and processes.

Here, we provide a synthetic overview of “sex in the wild” studies, the first of which we are aware since Bell (1982). We addressed this goal by performing a survey of directly relevant empirical literature to (1) assess the contributions of field studies towards resolving the sex problem, (2) identify what we have learned from these studies and what we still need to know, and (3) provide some concrete steps forward.

## Methods

### Literature survey approach

We began our survey by establishing a set of *a priori* criteria for study inclusion. These criteria were formulated with the goal of only including studies that could directly inform the maintenance of sex in natural populations. First of all, this meant that the study had to be performed in the field or use field-collected individuals that were not subsequently subject to the potential for selection in a laboratory environment. We included studies with laboratory-reared individuals only in those cases where lab rearing was unlikely to influence the factors that are the focus of testing (e.g., establishment of phylogenetic relationships). We therefore excluded studies where the potential for laboratory-imposed selection could confound the ability to interpret the study outcome, e.g. via laboratory-cultured lineages (e.g., Xu et al. 2011).

Second, we confined our survey to studies explicitly focused on investigating a particular mechanism for the maintenance of sex. We chose to adopt this mechanism-centered approach because this type of research takes place in a structured framework that facilitates meaningful comparisons among studies. Our strategy does have some limitations in excluding other valuable types of studies, including but not limited to evaluation of population genetic principles in natural populations (e.g., Menken et al. 1995, Lo et al. 2009), establishing the biogeographic distribution of sexual vs. asexual reproduction (e.g., geographic parthenogenesis; see Tilquin and Kokko 2016 for a recent overview), or describing unusual natural manifestations of reproductive strategies (e.g., Aanen et al. 2016).

Finally, we only included studies featuring obligately asexual individuals and facultatively or obligately sexual individuals that are sympatric in nature in at least part of their range. We made this choice because our goal is to identify the mechanisms that underlie the maintenance of sex in natural populations. Field-based studies in systems that feature sexual/asexual sympatry mean that the sexuals and asexuals can experience direct competition and allow for the most direct comparisons (with the fewest confounding factors) between reproductive modes. We do acknowledge that this particular approach does have limitations, such as the inability to directly address why some sexual lineages are not subject to invasion by asexual counterparts. We excluded all studies of sperm-dependent asexual taxa (e.g., *Ambystoma* mole salamanders, *Ips* bark beetles, *Rubus* subgenus *Rubus* blackberries) because all-asexual populations of sperm-dependent organisms are not evolutionarily stable.

We then classified and described each study according to the mechanism that the study addressed (as outlined below), the methods used, the study outcomes, whether the study supported or did not support the focal mechanism, and the taxonomic group represented by the focal organisms (see supplementary material; Table S1). We included an “Other” category to account for studies that addressed mechanisms or conditions with the potential to favor sex that did not easily fit under the umbrella of an established mechanism (e.g., reproductive assurance, which is largely confined to asexuals but needs to be tested by comparing sexuals and asexuals). We classified the species by class, following the U.S. Interagency Taxonomic Information System; when species from multiple classes were addressed, we used the lowest shared taxonomic level.

We included individual studies in more than one mechanism category if (a) the study found evidence for or against multiple mechanisms or (b) whether and how the study tested mechanisms was not clear enough to identify a single mechanism that was the focus of the paper. In the latter situation, we assessed each study carefully and then assigned the study to each mechanism that in our view was tested by the study design and/or was supported by the evidence delivered by the study. We termed each incidence of a distinct test of a distinct mechanism a “case”, which meant that some studies are represented by more than one case. We use this terminology consistently throughout the rest of the paper. We concluded our survey at the end of the summer of 2017, meaning that only papers published by this time were able to be included.

### Mechanisms underlying the benefits of sex

Sex, via segregation and recombination, breaks up linkage disequilibria (LD) across loci (Hill and Robertson 1966). This consequence of sexual reproduction is the reason that most mechanisms for the maintenance of sex focus on identifying conditions or situations associated with benefits of breaking up LD. Theoretical analyses have highlighted situations where selection changes over time and/or space or when linkage is generated by the combination of selection and drift as the conditions that are most likely to produce such benefits (Barton 2009). Field-based studies of sex that address benefits of LD breakup have typically focused on ecological situations (e.g., coevolving parasites or spatially structured niches, see below) that are expected to translate into changes in selection over space or time, but do not generally establish explicit links to the genetic mechanisms conferring benefits to sex. For this reason we *a priori* delineated the major mechanisms for sex by ecological mechanisms or scenarios that can generate or are associated with changes in selection.

We acknowledge several limitations of our approach to mechanism characterization.First, some of these categories feature conceptual overlap (e.g., “parasites” and “increased rate of adaptive evolution”, from the perspective that sexuals that can adapt more quickly to parasites might be at an advantage), while other categories could be combined to make even broader categories (e.g., all niche-based mechanisms; all hypotheses generated by genetic linkage).Second, it is virtually certain that some evolutionary biologists would produce different classification systems. Finally, we decided to separate mechanisms involving disadvantages to asexual lineages via reduced rates of adaptive evolution and increased rates of harmful mutation accumulation into separate categories. Although both phenomena are the consequence of the reduced efficacy of selection at linked sites (Hill-Robertson effect; Hill and Robertson 1966, Felsenstein 1974), different empirical methods are typically applied to detect evidence for ineffective adaptive evolution vs. ineffective purifying selection.

### Parasites

Biological antagonism –what we henceforth refer to as “parasites” or parasite pressure”– is potentially connected to the maintenance of sex because parasites can generate rapid changes in the direction of selection. The most prominent of the parasite mechanisms is the “Red Queen” (Bell 1982, Jaenike 1978, Hamilton 1980), under which parasites can help maintain sex by imposing negative frequency-dependent selection favoring rare host genotypes. Links between sex and biological antagonism can be driven by other mechanisms (e.g., Haafke et al. 2016), though the Red Queen-sex connection seems to be the most theoretically robust (e.g., Hamilton et al. 1990, Howard and Lively 1994, Peters and Lively 1999) and has received more empirical attention and support than other parasite-driven mechanisms for sex (reviewed in Neiman and Koskella 2009, Lively and Morran 2014).

### Rate of adaptive evolution

Selection works most effectively if beneficial and deleterious mutations occur in different individuals because these individuals should experience larger differentials in relative fitness than when mutations with opposite fitness effects co-occur within individuals (Hill and Robertson 1966). Over time, drift in the presence of selection may therefore lead to the accumulation of genomes where favorable and harmful alleles are linked. Sex can generate benefits in this situation in two ways: by immediately exposing ‘hidden variation’ (both favorable and harmful) to selection and by enabling more effective adaptive evolution at longer time scales.

### Harmful mutations

Hill-Robertson effects are expected to translate into increased rates of harmful mutation accumulation via relatively ineffective purifying selection in asexual vs. sexual lineages. Muller’s ratchet, which will cause irreversible mutation accumulation in small populations, is also expected to disproportionately affect asexual lineages (Muller 1964).

### Niche differentiation

One of the simplest mechanisms that enables coexistence between asexuals and their sexual relatives is niche differentiation. In the most extreme case of non-overlapping sexual and asexual niches, there is no competition between sexuals and asexuals, rendering costs of sex irrelevant (Meirmans et al. 2012). Perhaps because of its simplicity, there exist only a few theory-focused papers on this topic. One exception is the modelling study by Case and Taper (1986), who showed that niche differentiation can arise through character displacement after invasion of a sexual population by an asexual lineage. In practice, however, this mechanism is difficult to test: it is challenging to estimate the degree of niche overlap in natural populations and to determine whether the observed degree of niche differentiation between sexuals and asexuals is enough to prevent competition-driven extinction of one of the reproductive modes.

### Niche breadth

There are a variety of formulations of the overarching mechanism that sex can be maintained in situations where sexual individuals, lineages, or populations cover larger fractions of the available niche space than asexual counterparts. The most prominent example is the Tangled Bank hypothesis (Bell 1982), which postulates that sexual reproduction can be favored by generating a genetically diverse set of offspring that can make efficient use of a heterogeneous habitat via reduction in competition between siblings for limited resources. Asexually produced siblings, on the other hand, will compete for these same resources because they are genetically similar (also predicted by the conceptually similar frozen niche concept (Vrijenhoek 1979)). This advantage of sexual reproduction is increasingly offset as asexual lineage diversity increases, assuming that higher asexual diversity translates into more variable resource utilization by the asexual population.

## Results

Our literature survey of 66 studies (83 cases; some studies focused on multiple mechanisms) addressing the mechanisms underlying the maintenance of sex in the wild revealed some clear patterns (Table 1). First, there is a distinct majority of support (56 cases) vs. lack of support (27 cases), perhaps reflecting a publication bias towards positive results. The parasite and niche differentiation mechanisms predominated amongst the studies featuring positive results; we elaborate on these and other mechanism-specific patterns below. Our survey also clearly showed that some taxa have been the focus of far more investigation than others. The vast majority of studies involved animal systems (54 studies; 82%), which themselves were dominated by gastropods (23 studies), branchiopods (eight studies), reptiles (eight studies), and insects (six studies). The remaining 16% of the studies were in plants (11 studies; all Magnoliopsida/angiosperms) and fungi (one study). Some systems were particularly heavily represented. For example, the gastropod *Potamopyrgus antipodarum* was featured in 16 studies (see supplementary material, Table S1, for an overview). Other taxa common in our survey included the branchiopod *Daphnia pulex* (seven studies), the gastropod *Melanoides tuberculata* (four studies), and the angiosperm *Taraxacum officinale* (four studies). Only two studies applied a broad taxonomic approach, using comparisons of the ecologies of hundreds of sexual and asexual taxa (Ross et al. 2013, van der Kooi et al. 2017) to detect associations with, or consequences of, reproductive mode variation that apply across taxa.

**Table 1.** Summary of the results of our literature survey. We list the number of studies that provided support (green) or no support (red) for the different categories of mechanisms for sex, separated by taxonomic group. In total, 66 studies and 83 cases were included; studies that tested multiple mechanisms or included multiple taxonomic groups were counted as “cases” multiple times. Numbers in brackets after the name of the taxonomic group indicate the number of studies for each group.

| Class/clade | Parasites | Rate of adaptive evolution | Harmful Mutations | Niche differentiation | Niche breadth | Other |
| --- | --- | --- | --- | --- | --- | --- |
| Reptiles (8) | 2 / 1 | - | - | 2 / 2 | 0 / 2 | - |
| Insects (6) | 0 / 2 | - | 2 / 1 | - | 0 / 2 | - |
| Branchiopods (8) | 0 / 1 | - | 4 / 0 | 2 / 0 | 0 / 1 | 1 / 0 |
| Ostracods (4) | 0 / 1 | 2 / 0 | - | 2 / 1 | 0 / 2 | - |
| Arachnids (1) | - | - | 0 / 1 | - | - | - |
| Arthropods; general (1) | - | - | 0 / 1 | - | - | - |
| Gastropods (23) | 17 / 4 | 0 / 1 | 4 / 0 | 1 / 0 | 1 / 0 | 3 / 1 |
| Rotifers (2) | - | - | 1 / 1 | - | - | - |
| Anopla (1) | - | - | 1 / 0 | - | - | - |
| Magnoliopsida (11) | 2 / 0 | - | 2 / 1 | 5 / 1 | - | 1 / 0 |
| Fungi (1) | - | - | - | 1 / 0 | - | - |
| TOTAL | 21 / 9 | 2 / 1 | 14 / 5 | 13 / 4 | 1 / 7 | 5 / 1 |

### Parasites

Our search revealed 30 cases (representing 29 studies; one study with mixed results contributed two cases) that considered the maintenance of sex from the perspective of selection imposed via parasitism, by far the most frequently tested of the five main mechanism categories that we distinguished. Twenty of these cases reported results consistent with the expectations of situations where parasites are contributing to the maintenance of sex. Nearly all cases (27/30) focused at least in part on the Red Queen. A distinct majority of these Red Queen cases (15/27) involved the interaction between the snail *Potamopyrgus antipodarum* and the trematode parasite *Microphallus ‘*livelyi’. The three parasite-focused papers that did not explicitly address the Red Queen considered broader formulations of the parasite/antagonism mechanism (e.g., “parasitism rate”, “herbivory”).

### Rate of adaptive evolution

Our survey identified only three cases that addressed rates of adaptive evolution in the context of the maintenance of sex. All three of these cases took what we viewed as indirect approaches to this question, addressing whether sexuals were more often found in unpredictable abiotic environments that would likely demand rapid adaptation relative to the environments harboring asexual counterparts. Two ostracod-focused cases reported evidence consistent with this prediction, with sexual ostracods associated with relatively harsh and unpredictable environments (Schmit et al. 2013a,b). By contrast, parasites seemed to be a more likely explanation of patterns of distribution of sexual vs. asexual New Zealand freshwater snails than rapidly changing abiotic components of the environment (Lively 1987). Direct empirical evaluation of whether sexual organisms feature higher rates of adaptive evolution than their asexual relatives in natural populations clearly deserves future attention. Despite ample evidence for the importance of drift in natural populations (e.g., all studies supporting the ‘mutation accumulation’ mechanism), we found no field-based studies that investigated whether sex generates short-term benefits by exposing hidden genetic variation to selection.

### Mutation accumulation

The prediction that deleterious mutations accumulate more rapidly in asexual than sexual lineages has been tested in multiple taxa. Most cases (14 out of 19 cases; Table 1) were consistent with a scenario where asexuals accumulate deleterious mutations more rapidly than sexual lineages. Even so, nearly all (15/19) cases are based on a handful of genes, raising the question of the extent to which these results are likely to hold for the genome as a whole. Only four studies extended analyses of deleterious mutation accumulation to the genome scale (Hollister et al. 2015, Ament-Velásquez et al. 2016, Brandt et al. 2017, Lovell et al. 2017). Three of these studies (Hollister et al. 2015, Ament-Velásquez et al. 2016, Lovell et al. 2017) found that, as expected, asexuals had a higher load of deleterious mutations than sexuals. The fourth study (Brandt et al. 2017) found that *sexual* taxa experienced more mutation accumulation than asexual counterparts. This latter study focused on asexual lineages that are extremely old (tens of million years since derivation from sexual ancestors), suggesting that the absence of deleterious mutation accumulation may have contributed to the long-term persistence of these lineages in the absence of sex.

### Niche differentiation

We identified 17 cases that considered the maintenance of sex from the perspective of niche differentiation. Together with harmful mutations, these studies featured the broadest taxonomic support of the mechanism categories, representing six of the 11 taxonomic classes in Table 1. Tests for niche differentiation were most common in angiosperms, representing six of 17 cases. Thirteen of these 17 cases found at least some support for niche differentiation between sexuals and asexuals. Only three of these cases took the critical and perhaps most challenging additional step of determining whether the observed niche differentiation is enough to eliminate competition between sexuals and asexuals (O’Connell and Eckert 2001, Lehto and Haag 2010, Schmit et al. 2013a). All three cases did indeed suggest that niche differentiation between sexuals and asexuals was substantial enough that direct competition is unlikely, which should in turn resolve the problem of sex by rendering costs of sex irrelevant.

### Niche breadth

We identified eight cases where niche breadth was compared between asexual taxa and sexual counterparts, representing five taxonomic groups. In all but one case, the asexuals were characterized by broader niches than their sexual relatives. While most of these studies just focused on one or a few systems, the two studies that evaluated hundreds of species reported the same pattern (Ross et al. 2013, van der Kooi et al. 2017). Because the maintenance of sex via differential coverage of niche space requires that *sexuals* cover more niche space (and not, as found here, asexuals), our survey suggests that niche-breadth related mechanisms generally do not contribute to the maintenance of sex in the wild.

## Discussion

Perhaps most prominently, our survey revealed an important role for niche differentiation mechanisms in the maintenance of sex. This finding is also consistent with the many examples of geographical parthenogenesis and the observation that sexual and asexual individuals or lineages often differ in other key elements of their biology or ecology (e.g., hybrid status, polyploidy, production of resting eggs) (see also Meirmans et al. 2012). It is important to note that our survey revealed pervasive niche differentiation between sexuals and asexuals even in the exclusion of all studies focused on sexual and asexual taxa with non-overlapping ranges. While including these latter studies would perhaps have suggested an even stronger pattern of niche differentiation, without sympatric sexuals and asexuals, it is impossible to determine whether niche differentiation is a consequence of evolutionary divergence (character displacement following competition between sexual and asexual lineages) versus other factors such as biogeographic history (secondary contact). Evidence for the potential importance of niche differentiation from the field is striking because this mechanism has not otherwise received prominent attention in reviews on the benefits of sex (see Sharp and Otto (2016) for a recent example). We suspect that this inattention might be at least in part linked to the scarcity of theoretical studies on niche differentiation.

In contrast to niche differentiation, there was very little support for the related but distinct niche breadth mechanisms such as Tangled Bank (Bell 1982). While this type of mechanism has received quite a bit of theoretical attention (e.g., Bell 1982, Case and Taper 1986, Pound et al.2002), they seem to have very little empirical backing: in our survey, we found seven examples of negative results and only one example of support among the eight niche breadth-focused cases in our survey. This finding is in agreement with the general sentiment from earlier sex-focused research that revealed little evidence for Tangled Bank-like mechanisms (Ellstrand and Antonovics 1985, Burt and Bell 1987, but see Song et al 2011).

Another striking finding of our study was the absence of field studies focused on testing whether an increased rate of adaptive evolution might help favor sexual over asexual organisms. While there is substantial theoretical (Barton 1995, Burt 2000) and lab-based (e.g., Kaltz and Bell 2002, Goddard et al. 2005, McDonald et al. 2016) support for an important role for sexual reproduction in facilitating adaptive evolution, we only found three field-based studies that directly addressed whether sexuals have an adaptability advantage relative to asexual counterparts. One likely explanation for this distinct difference is that adaptability hypotheses are more difficult to rigorously test in the wild and are thus difficult to propose, fund, or execute. We also cannot exclude the possibility that publication bias (e.g., the challenges in publishing negative results) plays a role here or throughout our survey.

Most of the positive evidence in our survey comes from studies addressing the Red Queen hypothesis, a particular formulation of the parasite mechanism for sex. A distinct majority of these Red Queen-focused studies found evidence that parasite-host interactions contribute to the maintenance of sex. While the relatively large number and generally positive outcome of these studies might be taken as evidence that the Red Queen can provide a general explanation for the maintenance of sex in natural populations, the fact that most of these cases (15/ 27;∼55%) involve a single study system, *Potamopyrgus antipodarum*, means that this conclusion would be premature. The issues posed by dominance of a particular study system with respect to tests of a particular mechanism are highlighted by the fact that 15/15 cases in the *Potamopyrgus* system are consistent with the expectations of the Red Queen, compared to only five of 12 cases from other taxa (see Table S1 for more details). The problem of non-independence that arises from multiple tests of the same mechanism in the same system are heightened when, as for many sexual/asexual systems, there is only one transition to asexual reproduction, and thus, only one possible phylogenetically independent comparison. In general, our take-home message is that emergence of any mechanism as one that confers broad explanatory power for understanding sex in nature will require support from a diverse array of natural systems.

While many cases (12/17) provide at least some support for mutation accumulation mechanisms, most of the confirmatory evidence comes from only three study systems: *Daphnia, Potamopyrgus*, and *Campeloma*. All directly relevant genome-scale analyses of deleterious mutation accumulation of which we are aware have found considerable among-gene variation with respect to the rate of deleterious mutation accumulation in sexual and asexual lineages.Because of this among-gene variation, the inferences generated by the 15 cases that only included a handful of genes must be viewed with caution. Indeed, one of the four cases that did investigate mutation accumulation at the whole-genome scale in the wild found that, contrary to predictions, sexual taxa experienced more mutation accumulation (Brandt et al. 2017). Finally, although there appears to be some general support for deleterious mutation accumulation in asexual lineages, it is important to note that this mechanism is unlikely to explain the maintenance of sex. The reason for this caveat is that mutation accumulation generates a long-term disadvantage for asexuality, whereas short-term advantages are required for the maintenance of sex within populations (Maynard Smith 1978).

No single mechanism emerged as being important to the maintenance of sex across all natural systems included in our survey. This result provides indirect support for the idea that different mechanisms for sex might be important for different taxa (see also Neiman et al. 2017). While the notion that multiple mechanisms are of relevance to the maintenance of sex in nature would not surprise most of the researchers who study this topic, our survey results emphasize the importance of including a variety of taxa and considering multiple mechanisms when studying the maintenance of sex. Direct tests of the importance of this type of pluralism are conceptually simple but logistically challenging: the simultaneous study of multiple mechanisms across a diverse array of appropriate taxa in natural settings. The related but distinct point regarding the existence of a variety of different evolutionary “schools” concerning the maintenance of sex (Gouyon 1999, Meirmans and Strand 2010) also highlights the value of research on the same mechanisms or systems by multiple independent investigator groups.

### Outlook

Important steps forward for field-based research on the maintenance of sex will ideally come from several angles, which should perhaps most prominently include rigorous evaluation of the Red Queen hypothesis for sex in a diverse array of systems and direct tests for adaptive evolution (especially short-term effects) in natural settings. Especially insightful results with respect to the latter could be obtained from field experiments where sexuals and asexuals are transferred to (1) relatively harsh and/or unpredictable habitats, and (2) a relatively stable habitat, and where adaptation to the environment is tracked over multiple generations (see Lavanchy et al. 2016 for an example). Other useful tests could come from creative leveraging of unpredictable events (e.g., floods, fire, or even climate change) that might be expected to enable the detection of rapid adaption. Finally, researchers could evaluate whether sex generates short-term benefits in natural populations via the exposure of hidden genetic variation to selection by comparing genetic variation for fitness in naturally occurring and coexisting sexual and asexual organisms over multiple generations.

With respect to other mechanisms for the maintenance of sex, our study suggests that broader attention to niche differentiation mechanisms would likely turn out to be fruitful. In particular, the application of mesocosm approaches that reasonably reflect inherent natural complexity could bridge field and laboratory insights (e.g., Ganz and Ebert 2010). Throughout, we expect that an increasing availability of genomic resources for non-model taxa that have achieved prominence as the focus of “sex in the wild” studies (e.g., *Potamopyrgus antipodarum* (Bankers et al. 2017); *Taraxacum officinale*, E. Schranz pers. comm.) will provide a critical means of testing mechanisms for sex that make specific predictions regarding molecular evolution. We also believe that “pluralist” approaches that explicitly consider the possibility that different mechanisms might be important for different taxa or that separate mechanisms can operate simultaneously or in concert will provide key advances (Neiman et al. 2017). Finally, our study invites a formal comparison of outcomes of field versus laboratory studies that address the maintenance of sex. Evaluating whether lab studies consistently deliver similar vs. different outcomes than field studies with respect to particular mechanisms or taxa would be especially illuminating.

## Acknowledgements

We acknowledge Curt Lively for useful discussions on the history and development of the Red Queen hypothesis for sex and members of the Neiman lab as well as several anonymous reviewers and Associate Editor Britt Koskella for helpful comments on earlier versions of the MS. TS was supported by SNSF grant PP00P3 139013.

